# Characterizing representational shaping of individual motor and object representations after sequence learning

**DOI:** 10.1101/2025.09.02.673680

**Authors:** Nina Dolfen, Amir Tal, Wim Fias, Lila Davachi

**Affiliations:** Department of Psychology, Columbia University, New York, NY 10027; Department of Experimental Psychology, Ghent University, Ghent, Belgium; Psychology Department and Department of Cognitive and Brain Sciences, Hebrew University of Jerusalem, Jerusalem, Israel; Department of Clinical Research, Nathan Kline Institute for Psychiatric Research, Orangeburg, NY 10962

## Abstract

Learning temporal regularities between pairs of events has been shown to shape neural representations in the medial temporal lobe, but it remains unclear whether representational changes generalize across memory domains. Here we used fMRI and multivoxel pattern similarity analyses to examine representational shaping as a consequence of motor and object sequence learning. Across two sessions, participants incidentally learned a fixed four-element sequence of finger movements or visual objects. We compared the pattern of blood oxygen level-dependent activity evoked by each object and finger movement before and after sequence learning. Analyses were performed on bilateral ROIs, including primary motor cortex (M1), lateral occipital complex (LOC), premotor cortex (PMC), striatum (STR) and the hippocampus (HC). Behaviorally, participants successfully learned the sequence in both domains. At the neural level, motor representations became more differentiated with learning in M1, PMC, STR, and HC, but not in LOC. This global differentiation was not specific to the sequence condition and was also observed after repeated co-occurrence of movements in a pseudorandom order. Critically, the HC differentiated between motor representations in the learned sequence in a unidirectional predictive manner whereby movements became more differentiated from the preceding than from the following element in the sequence. In contrast, object representations remained stable across sequence repetitions in all brain regions, even though participants successfully learned their temporal order.

## Introduction

Many everyday activities consist of structured sequences (e.g., making a cup of coffee or typing in your computer password). Learning and remembering these sequences is necessary to perform tasks more efficiently, make predictions about the future and plan upcoming actions. This capacity, often referred to as sequence learning, has been studied extensively in both the motor (e.g., using the serial reaction time and sequential finger tapping tasks; for a review see Doyon et al., 2018) and non-motor domains (e.g., learning temporal regularities among objects, shapes or letters; for a review see Davachi and DuBrow, 2015). Historically, these domains have been considered distinct on both functional and anatomical grounds, so the literatures advanced largely in parallel. Yet both domains share a core computational demand: temporally organizing percepts and actions so that recall can proceed in the correct order.

In the motor domain, the acquisition of a new sequence of movements is initially characterized by large gains in speed and accuracy followed by smaller gains in performance during subsequent practice sessions (Doyon and Benali, 2005; Doyon and Ungerleider, 2002; Karni et al., 1998). These performance improvements are thought to be mediated by a process called binding or chunking (Graybiel, 2008; Lashley, 1951; Verwey, 1996), which refers to the temporal integration of individual motor responses into a single unit or motor chunk. At the behavioral level, chunking shapes sequence execution with typical slower responses at chunk boundaries (e.g., Kennerley et al., 2004; Sakai et al., 2003; Verwey et al., 2009; Verwey and Eikelboom, 2003) and by the emergence of correlations between response times, first locally between adjacent responses, and later between more distant elements (e.g., Beukema et al., 2019; Verstynen et al., 2012). Importantly, chunking is a slow process that requires extensive practice (Verstynen et al., 2012). It has therefore been suggested that the initial acquisition of a motor sequence relies on a much faster associative process that links successive motor responses and establishes first-order transitions (Beukema and Verstynen, 2018; Du and Clark, 2017). Such rapid associative binding has been discussed more broadly as a form of predictive processing, whereby the brain uses transitional probabilities to anticipate upcoming motor actions (Beukema and Verstynen, 2018).

Several studies have indicated that the fast detection of transitional probabilities may rely, at least in part, on the medial temporal lobe, particularly the hippocampus (Higuchi and Miyashita, 1996; Schapiro et al., 2014, 2012; Turk-Browne et al., 2010, 2009; see Davachi and DuBrow, 2015 for a review). The idea is that repeated exposure to temporal regularities strengthens a predictive code in the hippocampus (Bar, 2009) that carries information about the typical order of items in a sequence. One way to test this is to examine whether representations of individual sequence elements change across sequence repetitions, also referred to as representational shaping. For example, it has been demonstrated that firing rates of human hippocampal neurons at time *N* became more similar to those at time *N*+*1* after repeated viewing of the same movie clips, suggesting forward prediction of the next event (Paz et al., 2010). Likewise, using a statistical learning paradigm, repeated pairing of two fractals led to increased similarity of fMRI patterns in the hippocampus and medial temporal lobe cortex (Schapiro et al., 2012). Most regions in the medial temporal cortex exhibited bidirectional shaping, whereas subregions in the hippocampus represented regularities in a forward-looking predictive manner (Schapiro et al., 2012). These results indicate that the medial temporal lobe encodes item transitions by increasing the representational similarity of their members. Interestingly, accumulating evidence also suggest that the hippocampus contributes to motor sequence learning (Albouy et al., 2013, 2008; Döhring et al., 2017; Fernández-Seara et al., 2009; Jacobacci et al., 2020; Pinsard et al., 2019; Schapiro et al., 2019; Schendan et al., 2003) and represents information about the temporal order of movements in the learned sequence (Dolfen et al., 2024; Temudo et al., 2025). Accordingly, this raises the question of whether hippocampal associative and/or predictive processes that support learning in the non-motor domain extend to motor sequences, or whether such processes are instead supported by different regions, possibly operating in parallel. Animal studies point to the premotor cortex and the striatum as likely candidates (Bartolo et al., 2014; Crowe et al., 2014; Merchant et al., 2013; Mushiake et al., 1991). In addition, most prior work in the non-motor domain has focused on representational shaping as a consequence of associative pairing of two items, leaving open whether and how this generalizes to multi-element sequences.

Accordingly, we investigated whether predictive coding–like representational shaping of elementary sequence elements is a domain-general mechanism in the human brain, emerging after both motor and object sequence learning, and whether this process is supported by common or distinct brain regions across memory domains. To address this, each subject took part in two consecutive magnetic resonance imaging (MRI) sessions (one week apart), learning one sequence per session: either a sequence of four visually cued finger movements or a sequence of four visually presented objects. Subjects viewed each sequence 56 times over the course of the session, with an explicit test of sequence memory 24 hours after the scanning session. For the motor task, we additionally acquired reaction times and accuracy as a measure of online learning.

To investigate representational shaping of item representations, we compared the pattern of blood oxygen level-dependent (BOLD) activity evoked by each object and finger movement before and after sequence learning. To control for general effects of repeated co-occurrence of items, we also investigated changes after a control task, in which a different subset of items was presented repeatedly but in a pseudorandom order. Analyses were performed on five bilateral ROIs, including two low-level cortical regions involved in stimulus-specific processing (movements: primary motor cortex, M1; objects: lateral occipital complex, LOC) and three associative regions implicated in temporal order processing: premotor cortex (PMC) and striatum (STR) for movements, and the hippocampus (HC) for both domains. One possibility is that sequence learning leads to bidirectional associative binding, such that items within a sequence become associated and therefore more similar to each other, possibly leading to the strongest similarity increases between temporally adjacent elements. Another possibility (not mutually exclusive) is that sequence learning triggers forward predictive coding, whereby representations of each item (N) become tuned to anticipate the next item (N+1) in a unidirectional way. A third option is that regions come to represent the repeated co-occurrence of items but not in sequence specific manner. Here, we tested which of these accounts better characterizes representational shaping within motor and non-motor domains. Based on prior evidence, we expected representational shaping after sequence learning to emerge in associative regions (HC for motor and motor, PMC and/or STR for motor), but not in stimulus-selective low-level regions (M1, LOC).

## Methods

### Participants

Twenty-five healthy, young adults were recruited to participate in the study. All participants were screened for MRI eligibility prior to participation and again at the beginning of each study session. All participants were right-handed (Edinburgh Handedness questionnaire; Oldfield, 1971) and had corrected-to-normal vision. They did not report any current or previous neurological or psychiatric diseases and were free of medications. None of the participants were musicians. All participants reported normal sleep quality and quantity during the month prior to the study, as evaluated with the Pittsburgh Sleep Quality Index Buysse et al., 1989. Two participants were excluded from further analyses because the fMRI session was aborted prior to completion due to anxiety during scanning. A total of 23 participants (mean age: 24.36, range: 19-35, 17 females) were considered for the analyses. The study protocol was approved by the New York University Committee on Activities Involving Human Subjects. All volunteers gave written informed consent prior to the start of the study and received financial compensation upon completion of the study.

### General Procedure

The experimental timeline spanned two weeks, with participants completing one imaging session per week (7 days interval between sessions; see Figure 1A). At the beginning of the first imaging session, participants were assigned to either the motor or object condition with the assignment of task type (week 1 vs. week 2) counterbalanced across participants. During the motor session, participants were cued to perform series of visually guided finger movements during a serial reaction time task. During the object session, participants viewed series of visual objects while they performed a target detection task. Each MRI session consisted of three pre-encoding functional localizer runs (referred to as pre-encoding snapshots, Fig. 1B), eight encoding runs (referred to as encoding, Fig. 1B), and a post-encoding functional localizer run (referred to as post-encoding snapshot, Fig. 1B). During the first half of encoding (runs 1 to 4), participants performed a random control task in which cued responses/objects followed a pseudorandom order. Then, during the second half of encoding (runs 5 to 8) and unbeknownst to the participant a repeating fixed sequential pattern was introduced in order to induce sequence learning. Participants learned one sequence per imaging session: either a sequence of four cue-response associations (motor session; Fig. 1A) or four objects (object session, Fig. 1A). At the end of each fMRI session as well as 24 hours later, we assessed explicit knowledge about the temporal order of objects/movements within the repeating sequence.

**Figure 1.**
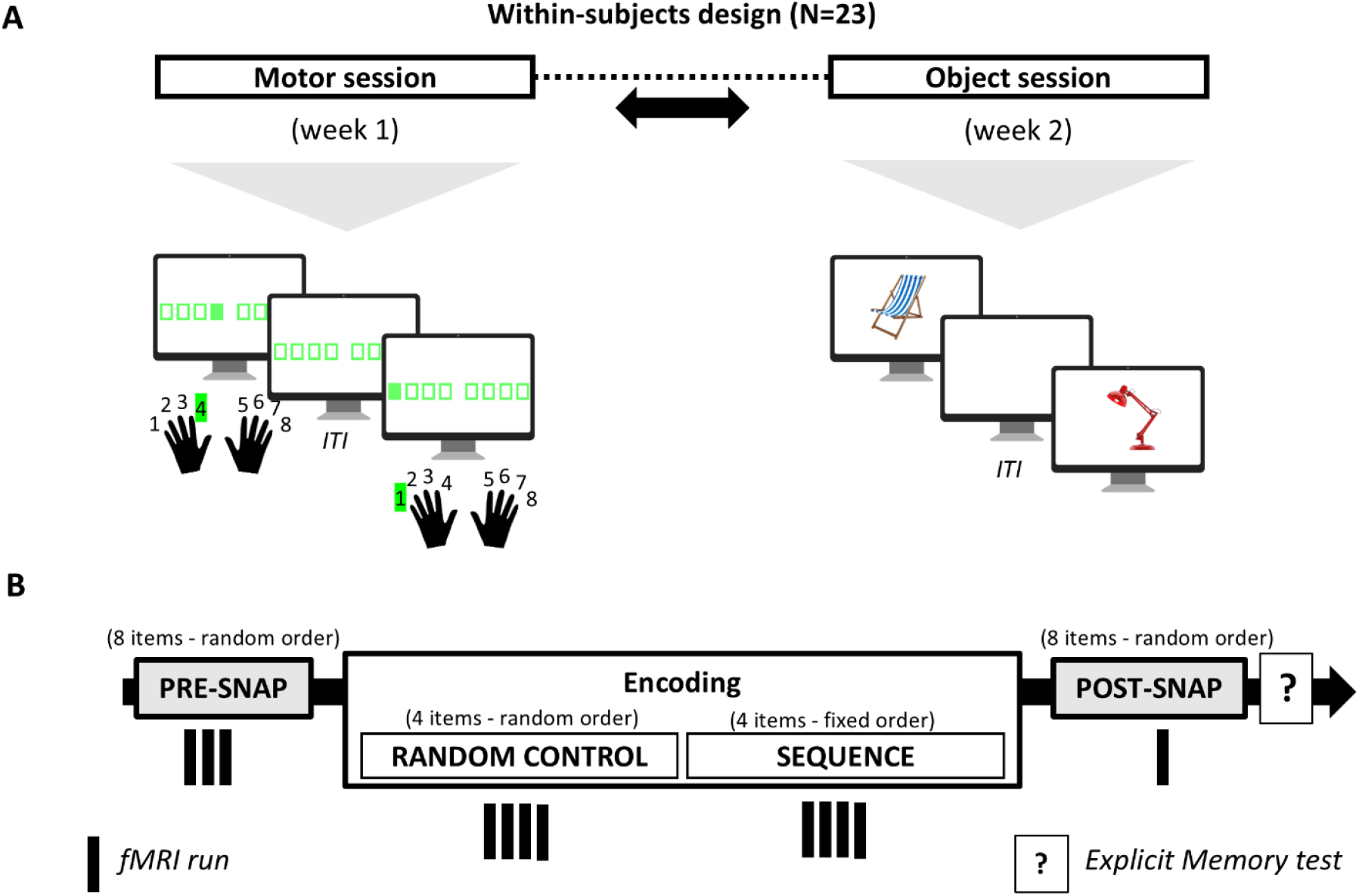
Study design and tasks. **(A)** Participants completed two imaging sessions spread across two weeks (with the order counterbalanced across participants). During the motor session, participants performed series of visually cued finger movements during a serial reaction time task (jittered inter-trial-interval; ISI; using 8 fingers, with 1 and 8 corresponding to the little fingers of the left and right hand, respectively, no thumbs). During the object session, participants viewed series of objects while they performed an orthogonal target detection task using their left or right index finger (jittered ITI; 8 different objects). **(B)** Each imaging session consisted of a pre-encoding localizer (i.e., pre-encoding snapshot, pre-snap), an encoding and a post-encoding localizer phase (i.e., post-encoding snapshot, post-snap). Of eight possible items (movements or objects) used in each session, four were assigned to the random control task and four to the sequence encoding task. During the random control task, items followed a pseudorandom order. During sequence encoding, items followed a fixed predetermined pattern to induce sequence learning. During the snapshots, all items were presented in a pseudorandom order to estimate the activation patterns evoked by each object (object session) or cued movement (motor session) before and after learning. The participants received no instructions on the pattern that the items might follow.

### Study design

During all imaging sessions, an event-related fMRI was used with jittered intervals between trials (see run specific sections for details). Each task involved eight items (8 cue-response associations during the motor session or 8 images of objects during the object session). During localizer runs, all eight items were included, whereas during encoding runs, half of the items were assigned to the random control condition and the remaining half to the sequence encoding condition. Hence, the subsets of items used in the random and sequence conditions were non-overlapping.

#### Pre- and post-encoding localizer runs

Participants completed three pre-encoding localizer runs and one post-encoding localizer run, referred to as snapshots. Data from the last pre-encoding run and the post-encoding run were used to estimate activation patterns evoked by each cue-response pair (motor session) or object (object session), before and after learning. During the localizer runs, items were presented in a pseudorandom order, with the constraint that no item repeated consecutively and each transition occurred equally often. The trial order in the last pre-encoding run was identical to the post-encoding run, allowing representational changes to be measured while minimizing concerns about temporal proximity artifacts. Each run included seven repetitions per item, with inter-trial intervals (ITIs) drawn from a truncated exponential distribution (1–5 s; TR-locked trial onsets).

#### Encoding

Encoding consisted of eight runs. In runs 1–4 (random control task), four items were presented in a pseudorandom order, while in runs 5–8 (sequence encoding task) the remaining four items followed a fixed, repeating sequence. Items appeared continuously with no explicit indication of sequence boundaries. The order in the random control task eliminated immediate repetitions to approximate the structure of the sequence encoding condition and each transition occurred equally often. Participants received no instructions about the presence of a sequence. Each run included 15 repetitions per item, with ITIs drawn from a uniform distribution (1.5–2.5 s; mean 2 s).

### Tasks

All tasks were implemented in Psychophysics Toolbox (Kleiner et al., 2007) in Matlab.

#### Motor tasks

During both localizer and encoding runs, participants performed a bimanual serial reaction time task. During the task, eight squares were presented on the screen, each corresponding to one of eight keys on the keyboard and one of eight fingers (corresponding to a total of 8 possible cue-response pairs, no thumbs; Fig. 1A). The outline of the squares alternated between red and green, indicating rest and practice, respectively. During rest, participants were instructed to keep their eyes focused on the screen and their fingers still on the keyboard (duration: 10s). During practice, the outline of the squares turned green, and participants were instructed to press as quickly and accurately as possible the key corresponding to the location of the cue (green filled square) that appeared on the screen. After a response, the next cue appeared with a jittered response to stimulus interval. Pre- and post-encoding snapshots included 56 trials each (7 repetitions × 8 cue-response pairs), while each encoding run included 60 trials (15 repetitions × 4 cue-response pairs). In addition, one 10 s rest periods was inserted halfway within each run. In the sequence encoding condition, participants were trained on one of two possible sequences of four finger movements (i.e. 8-3-5-2 or 1-6-4-7 with 1 and 8 corresponding to the little fingers of the left and right hand, respectively). However, to optimize the fMRI analysis, the starting point of the sequence was counterbalanced across participants so that the same key was not always associated to the same temporal position in the sequence. For each run (functional localizer and encoding runs), performance speed and accuracy were computed as the mean response time for correct responses (in seconds) and the percentage of correct responses, respectively. At the end of the session and again 24 hours later, participants completed a sequence generation task to assess explicit knowledge of the sequence. They were asked to self-generate, without visual cues, the sequence of finger movements performed during encoding (16 keypresses, corresponding to four repetitions of the four-element sequence).

#### Object tasks

During the object session, participants viewed series of visually presented objects while they performed an orthogonal target detection task. This ensured that participants maintained attention and processed the images. Stimuli consisted of eight colored object images (454 × 364 pixels). During the task, images were presented consecutively (stimulus duration: 1s; jittered interval between two images, see above). During pre- and post-encoding snapshots, participants were instructed to press a button (left or right) when a pound symbol appeared anywhere on the image (4% of trials; 7 x 8 objects = 56 trials per run). During encoding runs, participants performed a one-back task, pressing a button when an image repeated (5% of trials; 15 repetitions x 4 objects = 60 trials + 3 repetition trials per run). In the sequence encoding condition, participants repeatedly viewed the same sequence of four images (i.e. 1-2-3-4 or 5-6-7-8), with the starting point of the sequence varied across participants. For each run (functional localizer and encoding runs), accuracy (% correct hits) and reaction time for correct hits on the target detection task were measured. At the end of the session and again 24 hours later, participants performed a sequence generation task to assess sequence knowledge. They were shown the four sequence objects in random order and asked to self-generate the correct sequence, repeating this four times.

### Statistical analyses of behavioral data

For each task and measure, runs were excluded based on Tukey’s rule for outlier detection (1.5 × IQR; Tukey, 1977). To investigate behavioral changes during the motor session, we performed the following analyses: (1) one-way repeated-measure (RM) ANOVAs with run (4) as within-subject factor on speed and accuracy measures during the localizer runs and (2) condition (2) x run (4) RM ANOVAs on performance speed and accuracy during encoding. In the object session, we assessed whether target detection during localizer runs remained stable across localizer runs (one-way RM ANOVA) and compared target detection between encoding conditions (run x condition RM ANOVA). In case of violation of the sphericity assumption, Greenhouse–Geisser corrections were applied. To assess performance during the generation tasks, we computed the average proportion of correctly generated transitions across iterations as well as the proportion of times a full correct sequence was generated (maximum 4 times). For both measures we accounted for the possible rotations of the 4-item sequence (e.g., participants might have retained the sequence as 4-1-2-3 instead of 1-2-3-4). Performance was compared between time points and conditions using paired sample t-tests. Cohen’s d effect sizes were computed for t tests and partial eta squared (µ^2^) for F tests.

### MRI data acquisition

Images were acquired with a Siemens Prisma 3T MRI System and a 64-channel head coil. During the task runs, BOLD signal was acquired with a T2-weighted gradient-echo EPI sequence in axial orientation covering the whole brain (TR = 1500 ms; TE = 30 ms; FA = 65°; 69 slices; slice thickness = 2.0 mm; no interslice gap; FoV = 192 × 192 mm^2^; matrix size = 96 × 96 × 69; voxel size = 2.0 × 2.0 × 2.0 mm^3^; multiband factor = 3; GRAPPA = 2). A structural T1-weighted 3D MP-RAGE sequence (TR = 2.3s, TE = 2.98 ms, TI = 900 ms, FA = 9°, 176 slices, FoV = 256 x 248 mm^2^, matrix size = 256 × 248 × 176, voxel size = 1 × 1 × 1 mm^3^) and fieldmap (a single-echo EPI sequence original orientation: LAS, TR = 1.5 s, phase-encoding direction = posterior–anterior) were also obtained during each imaging session. The imaging session further contained three 6-min resting state fMRI scans. During the scans, participants viewed a blank gray screen and were instructed to remain awake while thinking about whatever they wanted. Analyses of the resting-state data are beyond the scope of this report. In total, each imaging session took about 90min to complete.

### MRI data preparation and preprocessing

Results included in this manuscript come from preprocessing performed using fMRIPrep 23.1.3 (Esteban et al., 2018), which is based on Nipype 1.8.6 (Gorgolewski et al., 2011). For more details of the pipeline, see the fMRIPrep documentation (https://fmriprep.org/en/stable/workflows.html). All study data were organized according to the Brain Imaging Data Structure (BIDS; Gorgolewski et al., 2016) specification, using *dcm2bids* (Boré et al., n.d.).

### Preprocessing of B0 inhomogeneity mappings

A *B*_*0*_ nonuniformity map (or *fieldmap*) was estimated from the phase-drift map(s) measure with two consecutive GRE (gradient-recalled echo) acquisitions. The corresponding phase-map(s) were phase-unwrapped with prelude (FSL None).

### Preprocessing of anatomical MRI data

A total of two T1-weighted images were part of BIDS dataset, one from each fMRI session. All of them were corrected were corrected for intensity non-uniformity (INU) with N4BiasFieldCorrection (Tustison et al., 2010), distributed with ANTs (Avants et al., 2008, RRID:SCR_004757), and used as T1w-reference throughout the workflow. The T1w-reference was then skull-stripped with a *Nipype* implementation of the antsBrainExtraction.sh workflow (from ANTs), using OASIS30ANTs as target template. Brain tissue segmentation of cerebrospinal fluid (CSF), white-matter (WM) and gray-matter (GM) was performed on the brain-extracted T1w using fast (FSL), RRID:SCR_002823, Zhang et al., 2001). Brain surfaces were reconstructed using recon-all (FreeSurfer 7.3.2, RRID:SCR_001847, Dale et al., 1999), and the brain mask estimated previously was refined with a custom variation of the method to reconcile ANTs-derived and FreeSurfer-derived segmentations of the cortical gray-matter of Mindboggle (RRID:SCR_002438, Klein et al., 2017). Volume-based spatial normalization to two standard spaces (MNI152NLin6Asym, MNI152NLin2009cAsym) was performed through nonlinear registration with antsRegistration (ANTs), using brain-extracted versions of both T1w reference and the T1w template. The following templates were selected for spatial normalization and accessed with *TemplateFlow* (23.0.0, Ciric et al., 2022): *FSL’s MNI ICBM 152 non-linear 6th Generation Asymmetric Average Brain Stereotaxic Registration Model* (RRID:SCR_002823; TemplateFlow ID: MNI152NLin6Asym), *ICBM 152 Nonlinear Asymmetrical template version 2009c* (Fonov et al., 2011, RRID:SCR_008796; TemplateFlow ID: MNI152NLin2009cAsym).

### Preprocessing of functional MRI data

For each BOLD run (across all tasks and sessions), the following preprocessing was performed. First, a reference volume and its skull-stripped version were generated using a custom methodology of *fMRIPrep*. Head-motion parameters with respect to the BOLD reference (transformation matrices, and six corresponding rotation and translation parameters) are estimated before any spatiotemporal filtering using mcflirt (FSL, Jenkinson et al., 2002). The estimated *fieldmap* was then aligned with rigid-registration to the target EPI (echo-planar imaging) reference run. The field coefficients were mapped on to the reference EPI. The BOLD reference was then co-registered to the T1w reference using bbregister (FreeSurfer) which implements boundary-based registration (Greve and Fischl, 2009). Co-registration was configured with six degrees of freedom. Several confounding time-series were calculated for each functional run based on the *preprocessed BOLD*: framewise displacement (FD), DVARS and three region-wise global signals. The three global signals are extracted within the CSF, the WM, and the whole-brain masks. Three probabilistic masks (CSF, WM and combined CSF+WM) were generated in anatomical space. Finally, these masks are resampled into BOLD space and binarized by thresholding at 0.99 (as in the original implementation, Behzadi et al., 2007). Following preprocessing using fMRIPrep, data in native space were used for further analyses.

### Regions of interest

All reported analyses were performed using a priori regions of interest. The following (bilateral) regions of interest (ROIs) were selected based on previous literature describing their involvement in stimulus-specific processes (object-selective or movement-selective) or in the processing of sequential information (Berlot et al., 2020; Davachi and DuBrow, 2015; Yokoi and Diedrichsen, 2019): primary motor cortex (M1), premotor cortex (PMC), lateral occipital complex (LOC), striatum (STR) and hippocampus (HC). Using Freesurfer’s automated cortical and subcortical segmentation (Fischl et al., 2002), we extracted each participants LOC, HC and STR (caudate and putamen). M1 and PMC were defined bilaterally in MNI space and next mapped back to native space using ANTs. Using the Brainnetome Atlas (Fan et al., 2016), M1 was defined to exclude mouth and leg representations and contained the upper limb and hand function regions of Brodmann area (BA) 4. Using the same atlas, the PMC was defined as the dorsal (A6cdl; dorsal PMC) and ventral (A6cvl; ventral PMC) part of BA 6. All ROIs were masked to exclude voxels outside of the brain.

### General linear modeling

Before conducting multi-voxel pattern analyses, we modeled the evoked response to individual cued movements and objects in the last pre-encoding as well as the post-encoding runs of the imaging session. For each participant, two separate GLMs were fitted to the preprocessed functional data (one for each session). The first GLM included one regressor per cue-response pair, separately for each timepoint (8 cue-response pairs × 2 timepoints = 16 regressors per run). The second GLM included one regressor for each object, separately for each timepoint (8 objects × 2 timepoints = 16 regressors per run). Within each regressor, events of interest were modeled with delta functions (0 ms for movements or 1 s duration for objects) time-locked to cue/object onsets. In both GLMs, incorrect responses and movement parameters (derived from realignment of the functional volumes; 6 motion parameters and their temporal derivatives) were modeled and entered as regressors of no interest. High-pass filtering with a cutoff period of 128 s served to remove low-frequency drifts from the time series. An autoregressive (Order 1) plus white noise model and a restricted maximum likelihood algorithm were used to estimate serial correlations in the fMRI signal. These GLMs generated separate maps of t values for each regressor in each run. Within each run, we removed voxels that were extremely inactive (Tambini and Davachi, 2013; Yu et al., 2024). To do so, separately for each participant, we computed the run-specific mean t-statistic across regressors for each voxel. We then calculated the mean and standard deviation of these values across all voxels within the mask and excluded voxels with a mean t-statistic more than two standard deviations below the mean. For each individual, the final t values (one value per voxel) were extracted and arranged into a vector per regressor, timepoint, and ROI. These vectors reflected the response of all voxels to each object and finger movement before and after encoding.

### Representational similarity analyses

RSA were designed to investigate changes in the representational dissimilarity between individual task elements (objects or movements) as a consequence of repeated encoding. Pattern dissimilarity was quantified as the Euclidean distance between multivoxel activity patterns evoked by different items before and after encoding. To prevent overemphasis of the population mean response dimension in fMRI voxels (Botero and Kriegeskorte, 2024), distances were computed after removing the mean of each pattern. For each subject, task, and timepoint, we computed two measures: (1) mean dissimilarity between all items in the learned sequence, and (2) mean dissimilarity between all items in the random control condition. Our primary analysis tested how representational distance changed from pre-to post-encoding, calculated as the difference in mean inter-item dissimilarity between timepoints separately for each encoding condition and task. To assess whether temporal proximity modulated representational change, we separately computed the change for consecutive (A–B, B–C, C–D, D–A) and non-consecutive transitions (A–C, B–D, C–A, D–B) in the learned sequence. Finally, to examine whether representational change reflected associative versus predictive processes, we assessed dissimilarity across timepoints. Specifically, we computed the distance between the post-encoding vector for item N and the pre-encoding vector for item N–1, and compared it with the reverse (post N–1 vs. pre N). A reliable difference between these values indicates asymmetric change (see results section).

To assess whether representational dissimilarity changed from pre-to post-encoding, a one-sample t test was performed for each ROI and encoding condition to compare average change to zero. Next, to compare representational changes (1) between different conditions (sequence vs. random control) or (2) between consecutive and non-consecutive items in the sequence, we used (two-tailed) paired sample t tests. Lastly, significant asymmetric changes within each ROI were tested with a one-sample t-test. For each test, Cohen’s d effect sizes were computed. For each group of hypotheses, the false discovery rate (FDR) procedure was applied to correct for multiple testing (i.e., corrected for 5 ROIs per test). Potential outliers were evaluated before statistical testing using Tukey’s rule (1.5 × IQR).

## Results

As illustrated in Figure 1A, all participants completed a motor and an object session. During the motor session, participants performed series of visually cued finger movements during a serial reaction time task. During the object session, participants viewed series of visually presented objects while they performed a target detection task. Within each session, items were presented under two encoding conditions (Fig. 1B): in the *random control* condition, items co-occurred in a pseudorandom order; in the *sequence* condition, items followed a fixed, repeating pattern. Of eight possible items (fingers/keys or objects) used per session, four were assigned to each encoding condition. Before and after encoding, all items were presented (objects) or executed (motor) again in a pseudorandom order (pre- and post-encoding snapshots, Fig. 1B), providing activation patterns for each item before and after learning (see fMRI methods section). This design allowed us to measure representational changes, while minimizing the risk that temporal proximity during encoding would artificially drive similarity between items (Schapiro et al., 2012).

### Behavioral results

#### Motor task

After the pre-encoding snapshot (see Supplementary Material for corresponding behavioral results), participants consecutively completed the random control and sequence encoding tasks. To investigate behavioral changes during motor encoding, a 2 (condition) x 4 (run) RM ANOVA was performed on both speed (reaction time) and accuracy (% correct responses). Behavioral results showed a condition by run interaction on both measures (RM ANOVA; interaction: speed, F_(3,66)_ = 3.42, *p* = 0.022; accuracy, F_(3,66)_ = 3.24, *p* = 0.027; main effect of block: speed, F_(3,66)_ = 15.48, *p* < 0.001; accuracy, F_(3,66)_ = 1.93, *p* = 0.133; main effect of condition: speed, F_(1,22)_ = 25.70, *p* < 0.001; accuracy, F_(1,22)_ = 0.93, *p* = 0.34). More specifically, while response times and accuracy remained stable during the random control task (simple main effect of run: speed, F_(3,66)_ = 1.61; µ^2^= 0.06; *p* =.194; accuracy, F_(3,66)_ = 0.20; µ^2^= 0.04; *p* =.894; Fig. 2A, control), both reaction times and accuracy improved during sequence encoding (simple main effect of run: speed, F_(3,66)_ = 18.06; µ^2^= 0.45; *p* <.001; accuracy, F_(3,66)_ = 6.40; µ^2^= 0.23; *p* <.001; Fig. 2A sequence). These results collectively suggest that participants learned the motor sequence. Explicit knowledge for the sequence was assessed with a sequence generation task, completed at the end of the imaging session (day 1) as well as 24hrs later (day 2). Participants recalled on average 39% of the response transitions (no difference between days, Fig. 2C) but were rarely able to generate the full sequence (i.e., only 26 % of participants reproduced the full sequence on day 1 and/or recalled it 24hrs later).

**Figure 2.**
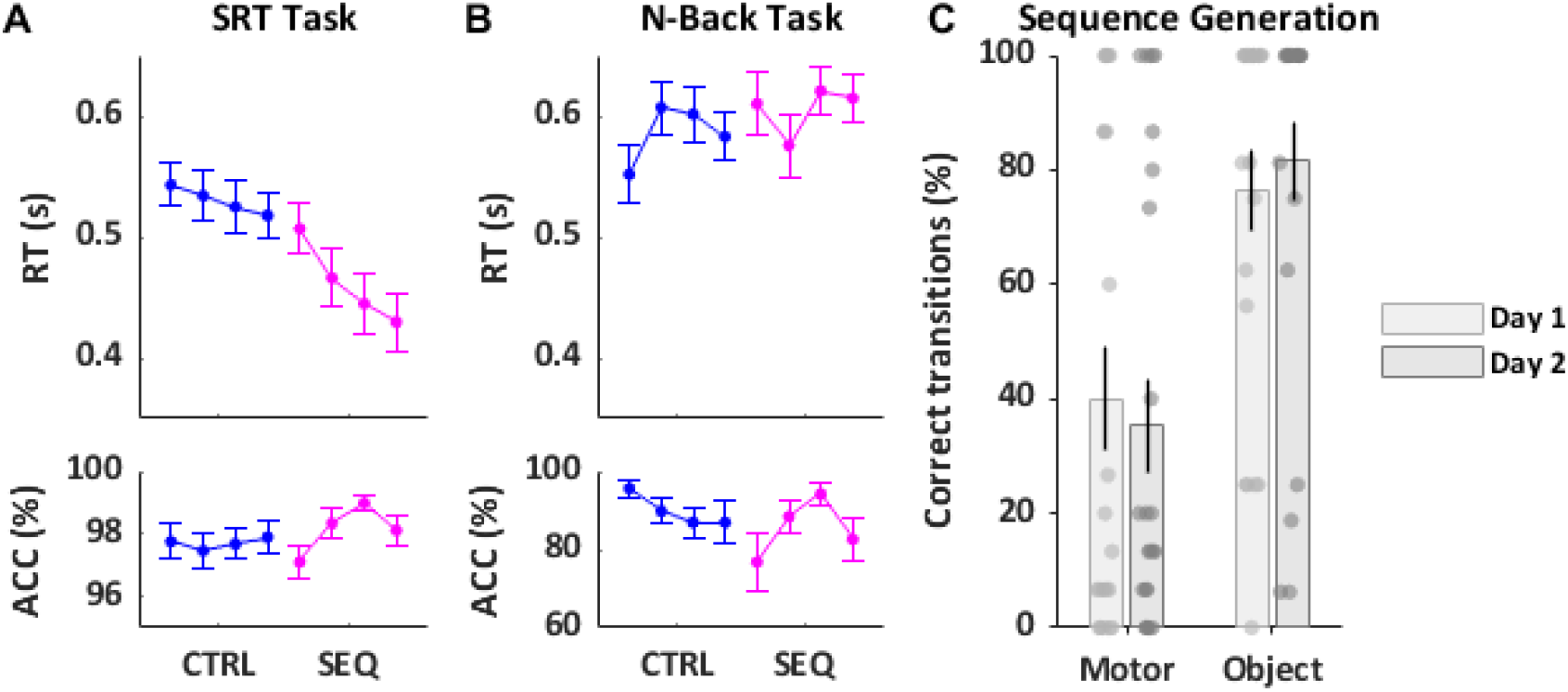
Behavioral results. **(A)** During motor encoding, participants performed a serial reaction time (SRT) task. Reaction time (RT; s) and accuracy (ACC; %) for the random control (CTRL) and sequence (SEQ) encoding conditions are plotted as a function of run in the fMRI session. **(B)** During object encoding, participants viewed series of objects while they performed an orthogonal 1-back task. RT (correct hits, s) and ACC (% correct hits) for the N-back task during the CTRL and SEQ encoding conditions are plotted as a function of run in the fMRI session. Note that in contrast to panel A, performance in panel B does not reflect whether participants learned the object sequence (but see panel B). **(C)** At the end of the imaging session (day 1) and 24hrs later (day 2), participants were tested on their sequence knowledge during an explicit sequence generation task. Percentages of correctly recalled item transitions are plotted as a function of session and day. N = 23. Error bars represent SEM.

#### Object task

During object encoding, reaction time on the target detection task remained stable across runs and conditions (condition x run RM ANOVA; main effect of run: F_(3,54)_ = 0.72, *p* =.543; main effect of condition: F_(1,18)_ = 1.47, *p* =.24; interaction: F_(3,54)_ = 1.84, *p* =.150). Average accuracy was not significantly different between conditions but there was a trend for a run x condition interaction (condition x run RM ANOVA; main effect of run: F_(3,66)_ = 0.89, *p* =.449; main effect of condition: F_(1,22)_ = 2.47, *p* =.130; interaction: F_(3,66)_ = 2.81, *p* =.05; Fig. 2B). This was driven by a marginally significant effect of run in the sequence encoding condition (simple main effect: F_(3,66)_= 2.22, *p* =.094), while no significant effect of run was observed in the random condition. Importantly, results of the sequence generation task indicated that participants acquired explicit knowledge about the temporal order of objects in the sequence during encoding. Participants correctly recalled on average 80% of the object transitions (no difference between days, Fig. 2C). When looking at the sequence level, results showed that 78.26% could reproduce the full sequence on day 1 and/or recall it 24hrs later.

### Neuroimaging results

We used multivoxel pattern similarity analyses of task-related fMRI data from the pre- and post-encoding snapshots to examine how individual movement or object representations are shaped by sequence learning. Specifically, we tested whether representational dissimilarity between items changed as a result of learning a structured sequence. To control for general effects of items repeatedly cooccurring together, we also assessed changes in representational dissimilarity for items in the random control conditions, where items were presented in a pseudorandom order. Analyses were performed on five bilateral ROIs: two stimulus-specific regions (M1 for movements; LOC for objects) and three associative regions implicated in temporal order processing (PMC, HC, STR).

#### Differentiation of motor representations but stable object representations

We first tested the *global representational change* after sequence learning, by assessing whether the overall dissimilarity between item representations (movements or objects) decreased from pre-to post-encoding. Such a decrease would indicate that activity patterns became *more similar* after sequence learning, consistent with associative binding. To test whether repeated co-occurence without sequential structure elicits similar changes, we performed the same analysis for items in the random control condition. For each ROI, we extracted the multi-voxel activity pattern elicited by individual finger movements or objects during the pre- and post-encoding snapshots, and then calculated, separately for each encoding condition, the change in average inter-item dissimilarity across timepoints. We next examined these global changes separately for the motor and object sessions.

#### Motor task

Contrary to our expectation of decreased dissimilarity, movement representations became more dissimilar over time. After sequence encoding, dissimilarity increased significantly in all regions except LOC (one sample t-test: all *p*_*corr*_ <.05; Fig. 3A; see caption for statistics). A similar pattern was observed after random control encoding, with significant increases in PMC, HC and STR (all *p*_*corr*_ <.05; Fig. 3A; see caption for statistics) but not in M1 or LOC. No significant differences were detected between sequence and random control conditions (paired sample t-tests, all *p*_*corr*_ >.1).

**Figure 3.**
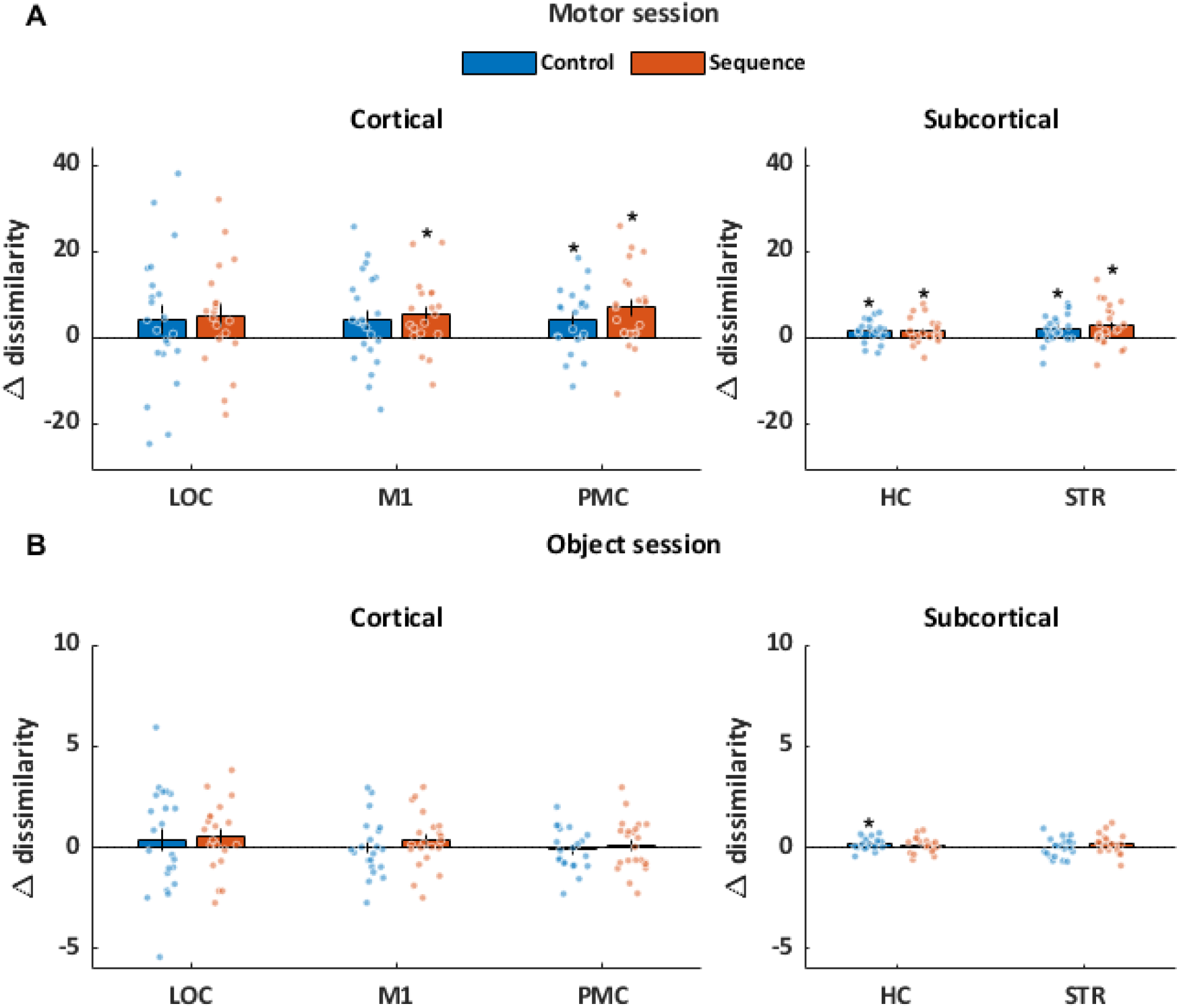
Global changes in representational dissimilarity (positive values indicate an increase) from before to after encoding as a function of encoding condition (random control vs. sequence) and session (motor vs. object). Motor representations became more dissimilar over time, whereas object representations remained largely stable. **(A)** After motor sequence encoding, dissimilarity increased in all regions (one sample t-tests: HC, t_(20)_ = 2.27, d =.49, *p*_*uncorr*_ =.034, *p*_*corr*_ <.05; STR, t_(22)_ = 2.82, d =.58, *p*_*uncorr*_ =.009, *p*_*corr*_ <.05; M1, t_(18)_ = 2.83, d =.65, *p*_*uncorr*_ =.011, *p*_*corr*_ <.05; PMC, t_(21)_ = 3.62, d=.77, *p*_*uncorr*_ =.0016, *p*_*corr*_ <.05) except for LOC (t_(19)_ = 1.84, d =.41, *p*_*uncorr*_ =.081). A similar pattern was observed after random control encoding, with significant increases in PMC (t_(21)_ = 2.61, d=.55, *p*_*uncorr*_ =.016, *p*_*corr*_ <.05), HC (t_(20)_ = 2.71, d =.59, *p*_*uncorr*_ =.013, *p*_*corr*_ <.05) and STR (t_(21)_ = 2.97, d =.63, *p*_*uncorr*_ =.007, *p*_*corr*_ <.05) but not M1 or LOC (both *p*_*cor*r_ >.1). **(B)** In the object session, representational dissimilarity was largely stable over time, with the exception of increased hippocampal dissimilarity after random control encoding (one-sample t-test: t_(19)_ = 2.89, d =.65, *p*_*uncorr*_ =.009, *p*_*corr*_ =.047), while all other effects were nonsignificant (all *p*_*corr*_ >.1). Abbreviations: M1, primary motor cortex. PMC, premotor cortex. LOC, lateral occipital complex. HC, hippocampus. STR, striatum. \**p*_*corr*_ < 0.05 based on one sample t-test. Error bars denote SEM.

#### Object task

In the object session, representational dissimilarity remained largely stable over time. The only reliable change was an increase in hippocampal dissimilarity after random control encoding (Fig. 3b; see caption for statistics). No significant changes were observed in other ROIs, nor were there differences between sequence and random control conditions (all *p*_*corr*_ >.1; Fig. 3B).

Taken together, these results show that motor representations became more differentiated with practice, regardless of sequence structure, whereas object representations were largely stable, with the exception of increased hippocampal dissimilarity for randomly presented items.

#### Representational shaping is not influenced by temporal proximity in the sequence

Although the global analyses above revealed increased differentiation for motor representations and largely stable object representations, representational shaping may be more localized — differently affecting consecutive as compared to non-consecutive items. Accordingly, we next compared average changes between consecutive (A–B, B–C, C–D, D–A) and non-consecutive transitions (all others) in both the motor and object sessions. However, we found no evidence that changes in representational dissimilarity differed between non-consecutive and consecutive items in either motor or object sequences (paired-sample t-tests, all *p*_corr_>.1, *p*_uncorr_ >.1; Fig. 4A, motor session; Fig. 4B, object session). Thus, we found no evidence for localized representational changes in either motor or object sequences.

**Figure 4.**
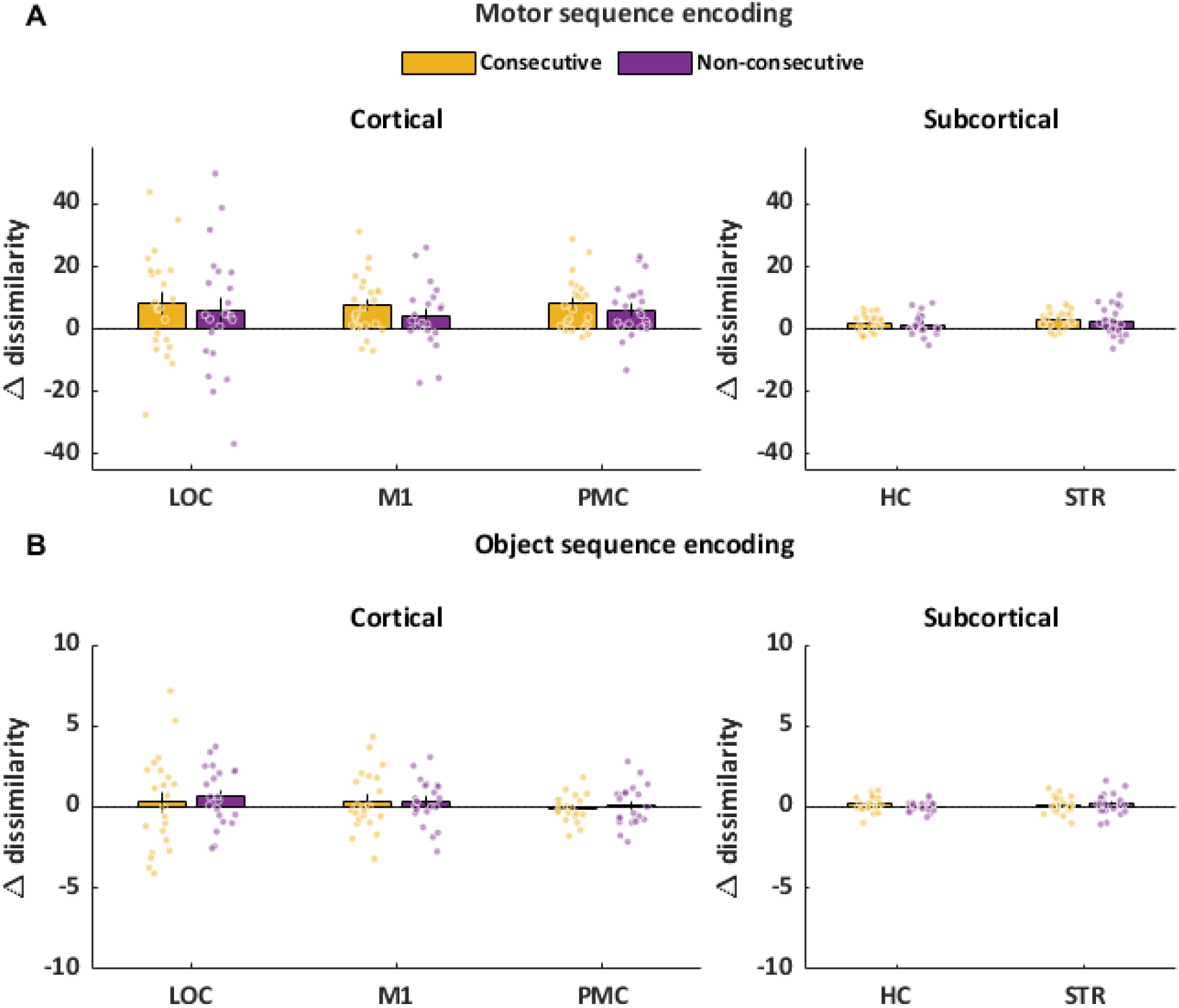
Changes in representational dissimilarity (positive values indicate an increase) from before to after sequence encoding as a function of temporal proximity in the learned sequence (consecutive vs. non-consecutive) for **(A)** the motor session and **(B)** the object session. Representational changes did not differ between consecutive and non-consecutive sequence elements in either session. SPL, superior parietal lobe. M1, primary motor cortex. M1, primary motor cortex. PMC, premotor cortex. LOC, lateral occipital complex. HC, hippocampus. STR, striatum. Error bars denote SEM.

#### Asymmetric differentiation of motor representations

So far, we have examined representational dissimilarity between items within the same timepoint, which only captures bidirectional changes. However, representational changes during sequence learning may be unidirectional, possibly revealing predictive mechanisms (Schapiro et al., 2012). To capture such asymmetries, we assessed dissimilarity across timepoints. Importantly, because asymmetric effects require a structured sequence and evidence for overall representational change, we restricted this analysis to ROIs in the motor sequence condition that showed a significant global change. Following prior work (Schapiro et al., 2012), we defined an asymmetry index between consecutive items N and N+1 (dissimilarity_(Npost, N+1pre)_ - dissimilarity_(N+1post,Npre)_), which will produce negative values if changes are backward-looking (i.e., N+1 became more dissimilar from the initial representation of the previous item N, than N from the initial representation of the subsequent item N+1) and positive values if they are forward-looking (i.e., N differentiated more from initial N+1, than vice versa). Across ROIs, only the hippocampus showed a reliable asymmetry (one sample t-test: t_(21)_ = 2.91, d =.65, *p*_*uncorr*_ =.008, p_corr_ <.05; Fig. 5), more specifically a negative asymmetry consistent with backward-looking differentiation from the preceding item. In all other regions, the asymmetry index did not differ significantly from zero, suggesting largely bidirectional changes. In this case, after learning, the representation of N became differentiated from how N+1 was initially represented, and this to the same extent as the representation of N+1 came dissimilar to how N was initially represented. As an exploratory follow-up, we asked whether the asymmetric change in the hippocampus was specific to the immediately preceding item. To this end, we compared dissimilarity between N_post_ and N–1_pre_ with dissimilarity to more distant items (N–2_pre_, N–3_pre_). Dissimilarity was significantly larger for lag –1 than for lags –2 or –3 (paired sample t-test; all p_uncorr_ <.05), while no difference was observed between lag –2 and lag –3, suggesting that the increased dissimilarity was specific to the immediately preceding item.

**Fig. 5.**
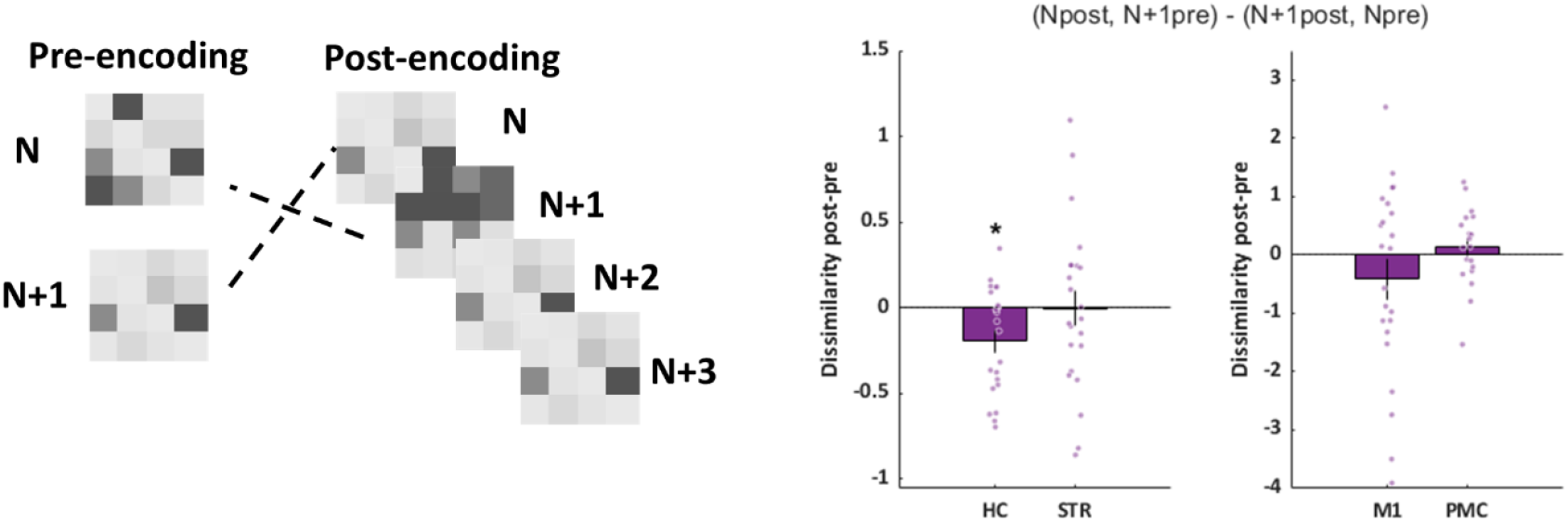
Asymmetric representational changes after motor sequence learning. **(A)** Schematic illustration of the across-timepoint comparisons. **(B)** Mean asymmetry index as a function of ROI with individual data points (dots). The asymmetry index (dissimilarity[Npost, N+1pre - dissimilarity[N+1post, Npre]]) was significantly negative in the hippocampus, consistent with backward-looking differentiation, but did not differ from zero in other ROIs, indicating bidirectional changes. Note that Y-axis scales differ between cortical and subcortical ROIs to account for differences in signal-to-noise ratio. M1, primary motor cortex; PMC, premotor cortex; LOC, lateral occipital complex; HC, hippocampus; STR, striatum. \**p*_*corr*_ < 0.05 based on one sample t-test. Error bars denote SEM.

## Discussion

In the current study, we used multivoxel pattern similarity analysis of fMRI data to investigate representational shaping of individual motor and object representations after motor and object sequence learning, respectively. To control for general effects of repeated co-occurrence, participants also completed a control task, in which a different subset of items were presented repeatedely but in a pseudorandom order. At the behavioral level, our data demonstrated that participants successfully acquired the motor and object sequences. At the neural level, we found evidence for increased differentiation of motor representations in M1, PMC, STR and HC post-as compared to pre-encoding. This global differentiation was not specific to the sequence condition, suggesting that repeated co-occurence of movements, whether in a structured or pseudorandom context, drives global differentiation between movement-specific fMRI patterns. Importantly, the hippocampus exhibited asymmetric differentiation of motor representations in the learned sequence, i.e., the post-encoding representation of item N+1 became more dissimilar from the pre-encoding representation of item N, but not vice versa. In the object session, object representations remained largely stable over time in all regions of interest and in both encoding conditions. Together, these findings indicate that, although both sequences were learned behaviorally, motor and object tasks had distinct effects on elementary neural representations.

Prior work in statistical learning has shown that the hippocampus can rapidly encode transitional probabilities between items in a task hippocampus (Higuchi and Miyashita, 1996; Schapiro et al., 2014, 2012; Turk-Browne et al., 2010, 2009). More specifically, Schapiro et al. (2012) demonstrated that repeated exposure to pairs of novel fractals led to increased pattern similarity between hippocampal representations of strongly associated items, but decreased similarity for weakly associated items, suggesting that the hippocampus is sensitive to learned probabilities by shaping individual item representations. Similarly, in the episodic memory domain (see Davachi and DuBrow, 2015 for a review), it has been shown that representational similarity in the hippocampus may be a mechanism for temporal memory organization (e.g., Ezzyat and Davachi, 2014). Motivated by this prior work and accumulating evidence for the role of the hippocampus in sequential processing across different memory domains (Döhring et al., 2017; Schapiro et al., 2019; Temudo et al., 2025), we asked whether the hippocampus might encode the temporal order of items during object and motor sequence learning by similarly shaping representations. Our data indicated that hippocampal representations of individual movements became more dissimilar, while no changes in hippocampal objects representations were detected after sequence learning. Critically, in the motor session, these hippocampal changes were asymmetric: movements became more differentiated from the preceding than from the following element in the sequence, revealing a backward-looking unidirectional change.

In the literature, differentiation between hippocampal representations after learning has been interpreted as a possible negative predictive code, whereby an item comes to predict the non-occurrence of another item (Schapiro et al., 2012). Given that the motor sequence in the current study was deterministic, this raises the question of why motor sequence learning led to backward-looking negative rather than forward looking positive prediction. One possibility is that the hippocampus supports sequence learning not only by linking items forward in time, but also by actively differentiating them from what has just occurred. Such backward-looking coding may serve to disambiguate adjacent events and maintain a clear temporal context, ensuring that each element is represented as distinct within the unfolding sequence. In this sense, the present finding could reflect a variant of negative prediction, with the hippocampus signaling that the prior event should not be reinstated once the next element has begun. In contrast to motor session, we found no evidence for representational changes in the object sequence. One possibility is that representations of well-known objects, rather than fractals which were used in previous work (Schapiro et al., 2012), are relatively robust and resistant to short-term plasticity, such that representational shaping of their associated representations may emerge only after more extended training. Thus, while prior studies have demonstrated predictive coding and associative shaping in the hippocampus during associative learning, the present data did not reveal such changes for object sequences.

Beyond the hippocampus, our analyses revealed increased differentiation between individual movement representations in M1, PMC, and STR after both random and sequence encoding. Importantly, the absence of change in this visual object-selective region suggests that differentiation is not a general property of all representations but emerges selectively in regions engaged in motor execution and/or temporal-order processing. The observation that task practice reshaped the representational distances in the motor cortical network contrasts with previous studies reporting stable motor representations after extended training on the serial reaction time task (Beukema et al., 2019). A key difference is that prior work examined changes across a 25-day training period, whereas the present analysis focused on a much shorter timescale within a single practice session. It is therefore possible that the differentiation observed here reflects more short-lived plasticity associated with initial learning rather than a more stable reorganization of motor representations once sequences are already learned and consolidated. This interpretation is reinforced by the fact that differentiation was also observed after the control task, which is less likely to induce durable plasticity (but see Ejaz et al., 2015 showing that activation patterns for finger movements in the primary motor and primary somatosensory cortices are determined by the natural, everyday statistics of how fingers move together). In summary, differentiation of motor representations in the cortico-striatal network may represent a temporary mechanism that supports accurate performance during the serial reaction time task.

Finally, we did not expect sequence-specific effects in M1 (Berlot et al., 2020; Yokoi et al., 2018; Yokoi and Diedrichsen, 2019), but we did hypothesize that the premotor cortex and striatum might support binding between movements during motor sequence learning. The absence of differences between random and sequence conditions is therefore unexpected. Both regions are strongly implicated in the acquisition and execution of action sequences (Graybiel, 2008; Hikosaka et al., 1999). The striatum, in particular, is thought to be important for the learning of predictive stimulus–response associations and is critical for motor chunking (Graybiel and Grafton, 2015; Penhune and Steele, 2012). Within this framework, striatal predictive processes in the current study may have been tied to anticipating the appropriate response from a given stimulus—an aspect that cannot be directly assessed with the present design, rather than encoding transitions between consecutive cued responses. Moreover, the striatum’s proposed role in motor chunking, a process that requires more extensive practice (Beukema and Verstynen, 2018), suggests that representational shaping of individual motor representations may emerge only after more extended training. This possibility is also consistent with its involvement in later stages of sequence automatization (Penhune and Steele, 2012). The premotor cortex, by contrast, has been linked to the representation of temporal features of motor sequences, such as the interval between key presses, irrespective of the specific movements performed (Kornysheva and Diedrichsen, 2014). Prior work also showed that premotor cortex activity increases when sequence structure becomes more complex, whether spatially or temporally (Bengtsson et al., 2004). In light of this, it is possible that our relatively simple four-element sequence was insufficient to elicit changes in the premotor cortex. However, given previous work, it seems more likely that sequence learning in the cortico-striatal network may not be supported by representational shaping at the level of individual sensorimotor representations, but rather by plasticity at a higher level of sequence or chunk representations (Berlot et al., 2020; Pinsard et al., 2019; Yokoi and Diedrichsen, 2019).

Before concluding, we offer some considerations related to the reported effects and the approach used in this study. First, the random control condition always preceded the sequence condition. This ordering was intentional, as an important goal was to test memory for the sequence after a consolidation interval. Had the random task followed the sequence task, it might have interfered with retention. At the same time, this design choice introduced a temporal confound that may have reduced sensitivity to sequence-specific effects. Second, the motor task may have been more engaging than the object task, potentially amplifying motor-related effects. Third, the analyses were based on relatively short training periods within a single session, which may capture transient plasticity but not longer-term representational reorganization.

In conclusion, the present study revealed overall differentiation between items in the motor domain, but no representational changes in the object domain. Importantly, hippocampal movement representations changed in an asymmetric, backward-looking manner, suggesting a novel mechanism for tracking temporal context. Whether this process is uniquely tied to motor sequence learning, or reflects a more general principle of hippocampal sequence coding, remains an open question for future work. In particular, given recent findings that temporal differentiation in event learning may be restricted to the dentate gyrus, with broader representational effects in CA3 (Bein and Davachi, 2024), future work employing high-resolution imaging could shed light on whether the asymmetries observed here reflect subfield-specific processes or a more general hippocampal principle.

## Notes

### Competing Interest Statement

The authors have declared no competing interest.

## Bibliography

Albouy G, King BR, Maquet P, Doyon J. 2013. Hippocampus and striatum: Dynamics and interaction during acquisition and sleep-related motor sequence memory consolidation. Hippocampus 23:985–1004. doi:10.1002/hipo.22183

Albouy G, Sterpenich V, Balteau E, Vandewalle G, Desseilles M, Dang-Vu T, Darsaud A, Ruby P, Luppi P-H, Degueldre C, Peigneux P, Luxen A, Maquet P. 2008. Both the Hippocampus and Striatum Are Involved in Consolidation of Motor Sequence Memory. Neuron 58:261–272. doi:10.1016/j.neuron.2008.02.008

Avants BB, Epstein CL, Grossman M, Gee JC. 2008. Symmetric diffeomorphic image registration with cross-correlation: Evaluating automated labeling of elderly and neurodegenerative brain. Med Image Anal 12:26– 41. doi:10.1016/j.media.2007.06.004

Bar M. 2009. The proactive brain: memory for predictions. Philosophical Transactions of the Royal Society B: Biological Sciences 364:1235–1243. doi:10.1098/RSTB.2008.0310

Bartolo R, Prado L, Merchant H. 2014. Information Processing in the Primate Basal Ganglia during Sensory-Guided and Internally Driven Rhythmic Tapping. Journal of Neuroscience 34:3910–3923. doi:10.1523/JNEUROSCI.2679-13.2014

Behzadi Y, Restom K, Liau J, Liu TT. 2007. A component based noise correction method (CompCor) for BOLD and perfusion based fMRI. Neuroimage 37:90–101. doi:10.1016/j.neuroimage.2007.04.042

Bein O, Davachi L. 2024. Event Integration and Temporal Differentiation: How Hierarchical Knowledge Emerges in Hippocampal Subfields through Learning. Journal of Neuroscience 44. doi:10.1523/JNEUROSCI.0627-23.2023

Berlot E, Popp NJ, Diedrichsen J. 2020. A critical re-evaluation of fmri signatures of motor sequence learning. Elife 9:1–24. doi:10.7554/eLife.55241

Beukema P, Diedrichsen J, Verstynen TD. 2019. Binding During Sequence Learning Does Not Alter Cortical Representations of Individual Actions. Journal of Neuroscience 39:6968–6977. doi:10.1523/JNEUROSCI.2669-18.2019

Beukema P, Verstynen T. 2018. Predicting and binding: interacting algorithms supporting the consolidation of sequential motor skills. Curr Opin Behav Sci 20:98–103. doi:10.1016/J.COBEHA.2017.11.014

Boré A, Guay S, Bedetti C, Meisler S, GuenTher N. n.d. Dcm2Bids. doi:10.5281/ZENODO.8436509

Botero VB, Kriegeskorte N. 2024. When do measured representational distances reflect the neural representational geometry? bioRxiv 2024.12.30.630743. doi:10.1101/2024.12.30.630743

Buysse DJ, Reynolds CF, Monk TH, Berman SR, Kupfer DJ, III CFR, Monk TH, Berman SR, Kupfer DJ. 1989. The Pittsburgh Sleep Quality Index: a new instrument for psychiatric practice and research. Psychiatry Res 28:193– 213. doi:10.1016/0165-1781(89)90047-4

Ciric R, Thompson WH, Lorenz R, Goncalves M, MacNicol EE, Markiewicz CJ, Halchenko YO, Ghosh SS, Gorgolewski KJ, Poldrack RA, Esteban O. 2022. TemplateFlow: FAIR-sharing of multi-scale, multi-species brain models. Nature Methods 2022 19:12 19:1568–1571. doi:10.1038/s41592-022-01681-2

Crowe DA, Zarco W, Bartolo R, Merchant H. 2014. Dynamic Representation of the Temporal and Sequential Structure of Rhythmic Movements in the Primate Medial Premotor Cortex. Journal of Neuroscience 34:11972–11983. doi:10.1523/JNEUROSCI.2177-14.2014

Dale AM, Fischl B, Sereno MI. 1999. Cortical surface-based analysis: I. Segmentation and surface reconstruction. Neuroimage 9:179–194. doi:10.1006/nimg.1998.0395

Davachi L, DuBrow S. 2015. How the hippocampus preserves order: The role of prediction and context. Trends Cogn Sci 19:92–99. doi:10.1016/j.tics.2014.12.004

Döhring J, Stoldt A, Witt K, Schönfeld R, Deuschl G, Born J, Bartsch T. 2017. Motor skill learning and offline-changes in TGA patients with acute hippocampal CA1 lesions. Cortex 89:156–168. doi:10.1016/j.cortex.2016.10.009

Dolfen N, Reverberi S, Beeck H Op de, King BR, Albouy G. 2024. The Hippocampus Represents Information about Movements in Their Temporal Position in a Learned Motor Sequence. Journal of Neuroscience 44. doi:10.1523/JNEUROSCI.0584-24.2024

Doyon J, Benali H. 2005. Reorganization and plasticity in the adult brain during learning of motor skills. Curr Opin Neurobiol 15:161–167. doi:10.1016/j.conb.2005.03.004

Doyon J, Gabitov E, Vahdat S, Lungu O, Boutin A. 2018. Current issues related to motor sequence learning in humans. Curr Opin Behav Sci 20:89–97. doi:10.1016/j.cobeha.2017.11.012

Doyon J, Ungerleider LG. 2002. Neuropsychology of Memory, Third Edition In: Squire LR, Schacter DL, editors. Neuropsychology of Memory, Third Edition. New York: Guilford Press. pp. 225–238.

Ejaz N, Hamada M, Diedrichsen J. 2015. Hand use predicts the structure of representations in sensorimotor cortex. Nat Neurosci 18:1034–1040. doi:10.1038/NN.4038

Esteban O, Markiewicz CJ, Blair RW, Moodie CA, Isik AI, Erramuzpe A, Kent JD, Goncalves M, DuPre E, Snyder M, Oya H, Ghosh SS, Wright J, Durnez J, Poldrack RA, Gorgolewski KJ. 2018. fMRIPrep: a robust preprocessing pipeline for functional MRI. Nature Methods 2018 16:1 16:111–116. doi:10.1038/s41592-018-0235-4

Ezzyat Y, Davachi L. 2014. Similarity breeds proximity: Pattern similarity within and across contexts is related to later mnemonic judgments of temporal proximity. Neuron 81:1179–1189. doi:10.1016/J.NEURON.2014.01.042/ATTACHMENT/ADEF83EB-1DBA-4EAA-9195-0DFD9346B9F1/MMC1.PDF

Fan L, Li H, Zhuo J, Zhang Y, Wang J, Chen L, Yang Z, Chu C, Xie S, Laird AR, Fox PT, Eickhoff SB, Yu C, Jiang T. 2016. The Human Brainnetome Atlas: A New Brain Atlas Based on Connectional Architecture. Cerebral Cortex 26:3508–3526. doi:10.1093/CERCOR/BHW157

Fernández-Seara MA, Aznárez-Sanado M, Mengual E, Loayza FR, Pastor MA. 2009. Continuous performance of a novel motor sequence leads to highly correlated striatal and hippocampal perfusion increases. Neuroimage 47:1797–1808. doi:10.1016/j.neuroimage.2009.05.061

Fischl B, Salat DH, Busa E, Albert M, Dieterich M, Haselgrove C, Van Der Kouwe A, Killiany R, Kennedy D, Klaveness S, Montillo A, Makris N, Rosen B, Dale AM. 2002. Whole Brain Segmentation: Automated Labeling of Neuroanatomical Structures in the Human Brain. Neuron 33:341–355. doi:10.1016/S0896-6273(02)00569-X

Fonov V, Evans AC, Botteron K, Almli CR, McKinstry RC, Collins DL. 2011. Unbiased average age-appropriate atlases for pediatric studies. Neuroimage 54:313–327. doi:10.1016/J.NEUROIMAGE.2010.07.033

Gorgolewski KJ, Auer T, Calhoun VD, Craddock RC, Das S, Duff EP, Flandin G, Ghosh SS, Glatard T, Halchenko YO, Handwerker DA, Hanke M, Keator D, Li X, Michael Z, Maumet C, Nichols BN, Nichols TE, Pellman J, Poline JB, Rokem A, Schaefer G, Sochat V, Triplett W, Turner JA, Varoquaux G, Poldrack RA. 2016. The brain imaging data structure, a format for organizing and describing outputs of neuroimaging experiments. Sci Data 3:1–9. doi:10.1038/SDATA.2016.44;SUBJMETA=114,2402,631,648,697,706;KWRD=DATA+PUBLICATION+AND+ARCHIVING,RESEARCH+DATA

Gorgolewski KJ, Burns CD, Madison C, Clark D, Halchenko YO, Waskom ML, Ghosh SS. 2011. Nipype: A flexible, lightweight and extensible neuroimaging data processing framework in Python. Front Neuroinform 5:12318. doi:10.3389/FNINF.2011.00013/BIBTEX

Graybiel A. 2008. Habits, rituals, and the evaluative brain. Annu Rev Neurosci 31:359–387. doi:10.1146/annurev.neuro.29.051605.112851

Graybiel AM, Grafton ST. 2015. The Striatum: Where Skills and Habits Meet. Cold Spring Harb Perspect Biol 7:a021691. doi:10.1101/CSHPERSPECT.A021691

Greve DN, Fischl B. 2009. Accurate and robust brain image alignment using boundary-based registration. Neuroimage 48:63–72. doi:10.1016/J.NEUROIMAGE.2009.06.060,

Higuchi SI, Miyashita Y. 1996. Formation of mnemonic neuronal responses to visual paired associates in inferotemporal cortex is impaired by perirhinal and entorhinal lesions. Proc Natl Acad Sci U S A 93:739–743. doi:10.1073/PNAS.93.2.739;REQUESTEDJOURNAL:JOURNAL:PNAS;CTYPE:STRING:JOURNAL

Hikosaka O, Nakahara H, Rand MK, Sakai K, Lu X, Nakamura K, Miyachi S, Doya K. 1999. Parallel neural networks for learning sequential procedures. Trends Neurosci 22:464–471. doi:10.1016/S0166-2236(99)01439-3

Jacobacci F, Armony JL, Yeffal A, Lerner G, Amaro E, Jovicich J, Doyon J, Della-Maggiore V. 2020. Rapid hippocampal plasticity supports motor sequence learning. Proc Natl Acad Sci U S A 117:23898–23903. doi:10.1073/pnas.2009576117

Jenkinson M, Bannister P, Brady M, Smith S. 2002. Improved optimization for the robust and accurate linear registration and motion correction of brain images. Neuroimage 17:825–841. doi:10.1006/nimg.2002.1132

Karni A, Meyer G, Rey-Hipolito C, Jezzard P, Adams MM, Turner R, Ungerleider LG. 1998. The acquisition of skilled motor performance: Fast and slow experience-driven changes in primary motor cortex. Proceedings of the National Academy of Sciences 95:861–868. doi:10.1073/pnas.95.3.861

Kennerley SW, Sakai K, Rushworth MFS. 2004. Organization of Action Sequences and the Role of the Pre-SMA. J Neurophysiol 91:978–993. doi:10.1152/JN.00651.2003

Klein A, Ghosh SS, Bao FS, Giard J, Häme Y, Stavsky E, Lee N, Rossa B, Reuter M, Chaibub Neto E, Keshavan A. 2017. Mindboggling morphometry of human brains. PLoS Comput Biol 13:e1005350. doi:10.1371/JOURNAL.PCBI.1005350

Kleiner M, Brainard DH, Pelli DG, Broussard C, Wolf T, Niehorster D. 2007. What’s new in Psychtoolbox-3? Perception 36:S14. doi:10.1068/v070821

Kornysheva K, Diedrichsen J. 2014. Human premotor areas parse sequences into their spatial and temporal features. Elife 3:e03043. doi:10.7554/eLife.03043

Lashley KS. 1951. The problem of serial order in behavior. New York: John Wiley Press.

Merchant H, Pérez O, Zarco W, Gámez J. 2013. Interval Tuning in the Primate Medial Premotor Cortex as a General Timing Mechanism. Journal of Neuroscience 33:9082–9096. doi:10.1523/JNEUROSCI.5513-12.2013

Mushiake H, Inase M, Tanji J. 1991. Neuronal activity in the primate premotor, supplementary, and precentral motor cortex during visually guided and internally determined sequential movements. 101152/jn1991663705 66:705–718. doi:10.1152/JN.1991.66.3.705

Oldfield RC. 1971. The assessment and analysis of handedness: The Edinburgh inventory. Neuropsychologia 9:97– 113. doi:10.1016/0028-3932(71)90067-4

Paz R, Gelbard-Sagiv H, Mukamel R, Harel M, Malach R, Fried I. 2010. A neural substrate in the human hippocampus for linking successive events. Proc Natl Acad Sci U S A 107:6046–6051. doi:10.1073/PNAS.0910834107/SUPPL_FILE/PNAS.200910834SI.PDF

Penhune V, Steele CJ. 2012. Parallel contributions of cerebellar, striatal and M1 mechanisms to motor sequence learning. Behavioural Brain Research 226:579–591. doi:10.1016/j.bbr.2011.09.044

Pinsard B, Boutin A, Gabitov E, Lungu O, Benali H, Doyon J. 2019. Consolidation alters motor sequence-specific distributed representations. Elife 8:1–20. doi:10.7554/eLife.39324

Sakai K, Kitaguchi K, Hikosaka O. 2003. Chunking during human visuomotor sequence learning. Exp Brain Res 152:229–242. doi:10.1007/s00221-003-1548-8

Schapiro AC, Gregory E, Landau B, McCloskey M, Turk-Browne NB. 2014. The Necessity of the Medial Temporal Lobe for Statistical Learning. J Cogn Neurosci 26:1736. doi:10.1162/JOCN_A_00578

Schapiro AC, Kustner L V., Turk-Browne NB. 2012. Shaping of object representations in the human medial temporal lobe based on temporal regularities. Current Biology 22:1622–1627. doi:10.1016/j.cub.2012.06.056

Schapiro AC, Reid AG, Morgan A, Manoach DS, Verfaellie M, Stickgold R. 2019. The hippocampus is necessary for the consolidation of a task that does not require the hippocampus for initial learning. Hippocampus 29:1091–1100. doi:10.1002/hipo.23101

Schendan H, Searl M, Melrose R. 2003. An FMRI study of the role of the medial temporal lobe in implicit and explicit sequence learning. Neuron 37:1013–1025. doi:10.1016/S0896-6273(03)00123-5

Tambini A, Davachi L. 2013. Persistence of hippocampal multivoxel patterns into postencoding rest is related to memory. Proc Natl Acad Sci U S A 110:19591–19596. doi:10.1073/pnas.1308499110

Temudo A, Dolfen N, King BR, Albouy G. 2025. The human medial temporal lobe represents memory items in their ordinal position in both declarative and motor memory domains. PLoS Biol 23:e3003267. doi:10.1371/JOURNAL.PBIO.3003267

Tukey JW. 1977. Exploratory Data Analysis. Reading, MA: Addison-Wesley. doi:10.1007/978-3-031-20719-8_2

Turk-Browne NB, Scholl BJ, Chun MM, Johnson MK. 2009. Neural evidence of statistical learning: Efficient detection of visual regularities without awareness. J Cogn Neurosci 21:1934–1945. doi:10.1162/JOCN.2009.21131

Turk-Browne NB, Scholl BJ, Johnson MK, Chun MM. 2010. Implicit Perceptual Anticipation Triggered by Statistical Learning. Journal of Neuroscience 30:11177–11187. doi:10.1523/JNEUROSCI.0858-10.2010

Tustison NJ, Avants BB, Cook PA, Zheng Y, Egan A, Yushkevich PA, Gee JC. 2010. N4ITK: Improved N3 bias correction. IEEE Trans Med Imaging 29:1310–1320. doi:10.1109/TMI.2010.2046908,

Verstynen T, Phillips J, Braun E, Workman B, Schunn C, Schneider W. 2012. Dynamic Sensorimotor Planning during Long-Term Sequence Learning: The Role of Variability, Response Chunking and Planning Errors. PLoS One 7. doi:10.1371/JOURNAL.PONE.0047336,

Verwey WB. 1996. Buffer Loading and Chunking in Sequential Keypressing. J Exp Psychol Hum Percept Perform 22:544–562. doi:10.1037/0096-1523.22.3.544

Verwey WB, Abrahamse EL, Jiménez L. 2009. Segmentation of short keying sequences does not spontaneously transfer to other sequences. Hum Mov Sci 28:348–361. doi:10.1016/j.humov.2008.10.004

Verwey WB, Eikelboom T. 2003. Evidence for lasting sequence segmentation in the discrete sequence-production task. J Mot Behav 35:171–181. doi:10.1080/00222890309602131,

Yokoi A, Arbuckle SA, Diedrichsen J. 2018. The role of human primary motor cortex in the production of skilled finger sequences. Journal of Neuroscience 38:1430–1442. doi:10.1523/JNEUROSCI.2798-17.2017

Yokoi A, Diedrichsen J. 2019. Neural Organization of Hierarchical Motor Sequence Representations in the Human Neocortex. Neuron 103:1178-1190.e7. doi:10.1016/j.neuron.2019.06.017

Yu W, Zadbood A, Chanales AJH, Davachi L. 2024. Repetition dynamically and rapidly increases cortical, but not hippocampal, offline reactivation. Proc Natl Acad Sci U S A 121:e2405929121. doi:10.1073/PNAS.2405929121/SUPPL_FILE/PNAS.2405929121.SAPP.PDF

Zhang Y, Brady M, Smith S. 2001. Segmentation of brain MR images through a hidden Markov random field model and the expectation-maximization algorithm. IEEE Trans Med Imaging 20:45–57. doi:10.1109/42.906424,

