## Supplementary Material for "Characterizing representational shaping of individual motor and object representations after sequence learning"

Figure S1, related to Figure 2

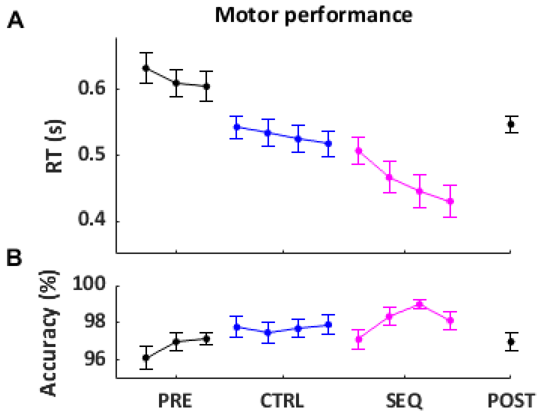


**Figure S1.** Behavioral results in the motor session. Reaction time (RT; s) and accuracy (ACC; %) for the pre-encoding localizers, random control and sequence encoding conditions as well as the post-encoding localizer are plotted across runs of practice in the fMRI session. N = 23. Error bars represent SEM. A RM ANOVA on motor performance indicated that speed (reaction time) increased (main effect of run: $F_{(3,66)}$ = 11.15; $\mu^2$= 0.34; $p$ < .001); while accuracy (% correct responses) remained stable across runs ($F_{(3,66)}$ = 0.95; $\mu^2$= 0.04; $p$ = .423; Fig. 3, snapshots). Follow-up pairwise comparisons indicated that reaction was significantly faster in the post-encoding snapshot as compared to the first and second pre-encoding run (run 1 vs. run 4, run 2 vs. run 4, both $p_{corr}$ < .05, all other comparisons, $p_{corr}$ > .05).
